# Scarcity of fixed carbon transfer in a model microbial phototroph-heterotroph interaction

**DOI:** 10.1101/2024.01.26.577492

**Authors:** Sunnyjoy Dupuis, Usha F. Lingappa, Xavier Mayali, Eve S. Sindermann, Jordan L. Chastain, Peter K. Weber, Rhona Stuart, Sabeeha S. Merchant

**Affiliations:** Department of Plant and Microbial Biology, University of California, Berkeley, CA 94720, USA; California Institute for Quantitative Biosciences (QB3), University of California, Berkeley, CA 94720, USA; Physical and Life Sciences Directorate, Lawrence Livermore National Laboratory, Livermore, CA 94550, USA; College of Chemistry, University of California, Berkeley, CA 94720, USA; Department of Molecular and Cell Biology, University of California, Berkeley, CA 94720, USA; Division of Environmental Genomics and Systems Biology, Lawrence Berkeley National Laboratory, Berkeley, CA, USA

**Keywords:** rhizobia, chlorophyte, photoautotroph, diel, cobalamin, co-cultivation, symbiosis, SIP

## Abstract

Although the green alga *Chlamydomonas reinhardtii* has long served as a reference organism, few studies have interrogated its role as a primary producer in microbial interactions. Here, we quantitatively investigated *C. reinhardtii’s* capacity to support a heterotrophic microbe using the established coculture system with *Mesorhizobium japonicum*, a vitamin B_12_-producing α-proteobacterium. Using stable isotope probing and nanoscale secondary ion mass spectrometry (nanoSIMS), we tracked the flow of photosynthetic fixed carbon and consequent bacterial biomass synthesis under continuous and diurnal light with single-cell resolution. We found that more ^13^C fixed by the alga was taken up by bacterial cells under continuous light, invalidating the hypothesis that the alga’s fermentative degradation of starch reserves during the night would boost *M. japonicum* heterotrophy. ^15^NH_4_ assimilation rates and changes in cell size revealed that *M. japonicum* cells reduced new biomass synthesis in coculture with the alga but continued to divide – a hallmark of nutrient limitation often referred to as reductive division. Despite this sign of starvation, the bacterium still synthesized vitamin B_12_ and supported the growth of a B_12_-dependent *C. reinhardtii* mutant. Finally, we showed that bacterial proliferation could be supported solely by the algal lysis that occurred in coculture, highlighting the role of necromass in carbon cycling. Collectively, these results reveal the scarcity of fixed carbon in this microbial trophic relationship (particularly under environmentally relevant light regimes), demonstrate B_12_ exchange even during bacterial starvation, and underscore the importance of quantitative approaches for assessing metabolic coupling in algal-bacterial interactions.

## INTRODUCTION

Photosynthetic microbes are responsible for a large fraction of the primary productivity on our planet [1]. These organisms supply the microbial world with raw materials for life, thereby shaping communities and driving carbon cycling [2, 3]. Through exudation, lysis, and predation, photosynthetic microbes provide a variety of reduced carbon substrates with various diffusion rates and accessibility to heterotrophic bacteria. The primary consumers recycle carbon back to inorganic forms, closing the “microbial loop” that constitutes a major component of biogeochemical carbon cycling [2, 4, 5]. Heterotrophic bacteria can also influence algal productivity, for example, by changing the availability of critical micronutrients for algal growth and metabolism [6–8].

The green alga *Chlamydomonas reinhardtii* has served as a reference organism for discoveries in photosynthesis, cilia-based motility, and the eukaryotic cell cycle [9]. Extensive work on axenic (bacteria-free) cultures has provided detailed knowledge of the pathways and regulons for photosynthesis and primary metabolism. This chlorophyte has a flexible metabolic repertoire, with the capacity to grow by photoautotrophy alone or by acetate assimilation. Under diurnal cycles, *C. reinhardtii*’s metabolism is shaped by programmed waves of gene expression that accommodate diurnal changes in light and oxygen availability in natural environments [10, 11]. Photosynthesis fuels cell growth and starch synthesis during the day, and starch reserves are glycolytically degraded at night, providing a source of energy until the next dawn. Although extensive knowledge of its metabolism makes *C. reinhardtii* an attractive model organism to study microbial trophic interactions, relatively little is known about how *C. reinhardtii* contributes to its environment and influences its neighbors in the microbial world.

In its natural soil context, *C. reinhardtii* is surrounded by other microbes that exchange nutrients, compete for resources, and communicate with one another. *C. reinhardtii* can enrich for particular heterotrophic bacteria from bulk soil, demonstrating that the alga’s growth can influence the metabolisms and behaviors of various bacteria [12, 13]. Studies of *Chlamydomonas*-bacteria interactions have primarily focused on impacts to the alga’s behavior, in part due to technical challenges in probing bacterial physiology in mixed cultures. Bacteria are known to supply vitamins and fixed nitrogen, facilitate iron assimilation, and impart thermal tolerance to algal cultures [6, 7, 14–20]. Coculture of *C. reinhardtii* with various heterotrophic bacteria (e.g. *Bradyrhizobium japonicum*, *Pseudomonas* spp.) has been shown to stimulate algal H_2_ production, starch accumulation, and viability under sulfur deprivation by consuming O_2_ and alleviating accumulation of fermentation byproducts from the medium [21]. Bacterial secondary metabolites can also influence *C. reinhardtii*’s cell cycle and aggregation [22, 23]. In addition, interactions between *C. reinhardtii* and vitamin B_12_-producing rhizobia like *Mesorhizobium japonicum* (previously referred to as *Mesorhizobium loti*), *Sinorhizobium meliloti*, and *Rhizobium leguminosarum* have been used to study the evolution of vitamin dependency across the algal tree of life [6, 15, 24–28]*. C. reinhardtii* does not require vitamin B_12_ for growth, because it has retained a functional cobalamin-independent isoform of methionine synthase, METE1. However, when vitamin B_12_ is supplied, *METE1* is repressed, and the vitamin is used as a cofactor for the cobalamin-dependent isoform, METH1. When *C. reinhardtii* was grown in the laboratory with a continuous supply of the vitamin, the alga acquired deleterious mutations in the repressed *METE1* gene, yielding B_12_-requiring strains (*mete1*) in less than 500 generations [25].

The interaction between the *C. reinhardtii mete1* mutant and *M. japonicum* is the most well-studied *C. reinhardtii* coculture system. It has been suggested to be facultatively mutualistic: the mutant alga receives the vitamin cofactor, and the bacterium is believed to receive reduced carbon compounds that support heterotrophic growth. However, the mechanism and magnitude of the flux of carbon transfer from *C. reinhardtii* to *M. japonicum* are unknown. Quantitative investigations of algal-bacterial interactions have been challenging due to technical barriers for studying eukaryotes and prokaryotes simultaneously. We have addressed this challenge by using spatiotemporal stable isotope tracing via nanoscale secondary ion mass spectrometry (nanoSIMS) to probe this simple model interaction – a technique which has typically been used for studying complex polycultures from the field. We quantified the transfer of fixed carbon from wild-type *C. reinhardtii* to *M. japonicum* in continuous and diurnal light. Although some photosynthetically fixed carbon was taken up by the bacterial population, *M. japonicum* growth was severely limited, resulting in reductive bacterial division. Thus, *C. reinhardtii* primary productivity yields a low carrying capacity for this bacterium, particularly under diurnal light. Nonetheless, *M. japonicum* provided adequate B_12_ to enable growth of the *C. reinhardtii mete1* strain even when the bacterium itself exhibited signs of starvation. These results highlight that in natural contexts, bacteria may not proliferate to the high cell densities that are possible in nutrient-rich media often used in the laboratory, but they can still greatly influence the metabolism and evolution of their neighbors in the microbial world.

## MATERIALS AND METHODS

### Strains and culture conditions

Three strains of *C. reinhardtii* were used in this study. The *mete1* strain generated through experimental evolution on 1000 ng/l vitamin B_12_ [25] and its parental wild-type strain were provided by Alison G. Smith (University of Cambridge, UK). According to its haplotype, this strain is closely related to strain S24-(SI Fig. 1) [29]. For experiments testing the impact of diurnal light, we used the cell-wall reduced strain CC-5390. This strain is amenable to synchronization under diurnal cycles, providing a system with exceptional signal-to-noise for resolving diurnal patterns [10]. *M. japonicum* strain MAFF303099 was also provided by Alison G. Smith.

**Figure 1:**
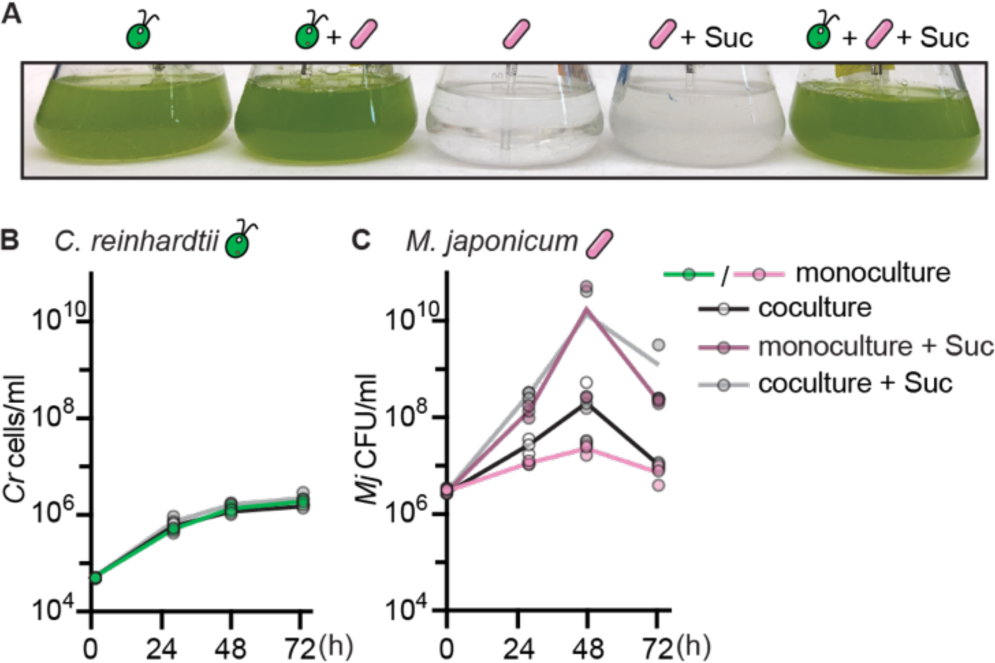
Wild-type *C. reinhardtii* does not significantly improve growth of *M. japonicum*. Triplicate cultures (*n* = 3) were inoculated with or without 150 µg/ml sucrose and maintained under continuous light, bubbled with air, and shaken at 110 rpm. (A) Representative image of cultures 72 h after inoculation. (B) Wild-type *C. reinhardtii* cell density in cultures over time after inoculation. (C) *M. japonicum* cell density in cultures over time after inoculation. The maximum bacterial density in coculture was not significantly different from the maximum bacterial density in monoculture with or without sucrose by two-tailed Student’s *t*-tests (*p* > 0.05).

All experiments were performed in amended HSM minimal medium [30] with a modified trace element solution [31], 2 mM MgSO_4_·7H_2_O, 10 µg/l biotin, and 10 µM CoCl_2_ unless otherwise specified (Supplementary Data File S2: Media composition). Cultures were grown in either 250 ml Erlenmeyer flasks containing 100 ml medium and bubbled with filter-sterilized air (provided by an aquarium pump), 6-well plates with 3 ml medium, or 250 ml beakers with 50 ml medium. Algal cultures and cocultures were incubated at 28°C, agitated at 110–130 rpm, and illuminated with 200 µmol photons/m^2^/s cool white light (Sylvania Dulux L55WFT55DL/841) continuously or under diurnal 12-h-light/12-h-dark cycles.

*C. reinhardtii* cells were precultivated axenically in photoautotrophic conditions for at least 3 passages prior to experiments. The *mete1* strain was precultivated with 200–250 ng/l vitamin B_12_ (cyanocobalamin). For experiments comparing coculture in continuous light versus diurnal light, *C. reinhardtii* CC-5390 cells were precultivated in flat-panel photobioreactors (Photon System Instruments, Drásov, Czechia) as previously described under the respective light regime to achieve adequate synchrony of the diurnal populations [10].

Bacterial cells were precultivated in 14 ml polystyrene round-bottom tubes with 3 ml amended HSM containing 0.2% sucrose, agitated at 200 rpm, and incubated at 28°C. When larger volumes were required for inoculation, cells were precultivated instead in 250 ml Erlenmeyer flasks containing 100 ml medium, continuously bubbled with air, and agitated at 110–130 rpm. Bacterial cells were collected by centrifugation at 8000 x*g* and washed three times in 0.85% NaCl to remove sucrose before use as inoculum. Optical density at 600 nm (OD_600_) was used to estimate the cell density of washed cell suspensions used for inoculation.

### Cell number, size, and cytotoxicity measurements

*C. reinhardtii* cell density and size were determined using a hemocytometer and ImageJ or a Beckman Coulter Multisizer 3 with a 50 μm orifice (Beckman Coulter, CA, USA). Bacterial cell density was determined by counting colony forming units (CFU) in 10 µl spots of serial dilutions on Typtone Yeast (TY) agar medium (Supplementary Data File S2: Media composition) after 4 d of incubation at 30°C in the dark.

Algal cellular integrity was estimated using CellTox Green Cytotoxicity Assay (Promega, WI, USA). *C. reinhardtii* sample densities were adjusted to 1×10^6^ cells/ml. Triplicate killed control samples were prepared by heating cells at 90°C for 10 min. Fluorescence was measured in a black-walled 96-well plate using a SpectraMax iD3 plate reader (Molecular Devices, CA, USA). Background fluorescence of a medium blank was subtracted from all values. Results were reported relative to killed control samples.

Samples for fluorescence microscopy were fixed overnight in 4% paraformaldehyde at 4°C in the dark. Fixed samples were then collected by centrifugation at 10,000 x*g*, washed in phosphate buffered saline (PBS) (pH 7), and stored in 1:1 ethanol:PBS at −20°C. For imaging, samples were spotted on 0.22 µm GTTP type isopore polycarbonate filters and washed with sterile water to remove residual salts. Prepared filters were mounted on slides with 5 µg/ml DAPI in Citifluor antifade mounting medium buffered with PBS and imaged using a Zeiss Axioimager M2 microscope with a DAPI filter set (Ex 350/50, Em 460/50). Bacterial cell lengths were measured manually using ImageJ Version 2.1.0/1.53c by drawing a line from one pole of the cell to the other along the longer axis.

### Total organic carbon measurements

Total nonpurgeable organic carbon content (NPOC) of cells was determined using a TOC-L Shimadzu Total Organic Carbon Analyzer (Shimadzu, Kyoto, Japan). 10–40 ml culture was collected by centrifugation at 8,000 x*g* for 2 min, and the supernatant was collected and then stored at −20°C until analysis. Spent media were diluted 2-fold and HCl was added to 27 mM. All samples were sparged to remove inorganic carbon. Some organic carbon may be purged from the sample by this method, so we report “nonpurgeable” organic carbon.

### Stable isotope labelling and nanoSIMS

^15^NH_4_Cl (99% ^15^N, Cambridge Isotope Laboratories, MA, USA) was provided directly in the culture medium, replacing 50% of the NH_4_Cl in HSM. 13 h after inoculation, ^13^CO_2_ was provided to cultures by bubbling air through a solution of NaH^13^CO_3_ (99% ^13^C, Cambridge Isotope Laboratories) and then into the cultures as previously described [32]. Killed controls were generated by treating cells with 4% paraformaldehyde for 20 min in the dark. Coculture experiments were otherwise conducted as described above.

For nanoSIMS analyses, samples were fixed and deposited on filters as described above for fluorescence microscopy. Dry filter pieces were then mounted on 1 inch round stubs using carbon tabs (Ted Pella, Inc., CA, USA) and gold coated. Data were collected on a NanoSIMS 50 (CAMECA, Gennevilliers, France) using a 1.5 pA Cs^+^ primary ion beam. The secondary ion masses of ^12^C_2_^−^, ^12^C^13^C^−^, ^12^C^14^N^−^, ^12^C^15^N^−^, and ^32^S^−^ were collected. Raw nanoSIMS images were processed using L’image (L. Nittler, Carnegie Institution of Washington, D.C., USA). Individual cell regions of interest (ROIs) were manually circled using size and high ^12^C^14^N signal to identify bacterial cells and high ^12^C^13^C signal along with the secondary electron image to identify algal cells. Isotope enrichment in atom percent enrichment (APE) for ^12^C^13^C^−^/^12^C_2_^−^ and ^12^C^15^N^−^/^12^C^14^N^−^ were calculated using standard ratios of 0.02247 and 0.00367, respectively. Enrichment data are also presented as net assimilation (N_net_ and C_net_). The fraction of newly synthesized biomass relative to total biomass was calculated against an unlabeled biomass standard as previously described [33]:

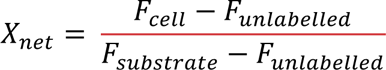

Where *F* is the abundance of the heavy isotope (^15^N or ^13^C) over the total (^14^N+^15^N or ^12^C+^13^C, respectively). For ^15^N, net assimilation was calculated relative to the known isotopic composition of the initial source substrate added to the medium. For ^13^C, net bacterial assimilation was calculated relative to the average algal biomass isotopic composition in the respective experiment (continuous light or diurnal light) as a rough estimation of the source substrate.

### Vitamin B_12_ bioassay

Vitamin B_12_ concentration was determined using the *Escherichia coli* B_12_ bioassay described previously [34, 35]. The bioassay uses two *E. coli* strains: the Δ*metE*Δ*metH* control strain requires methionine for growth, and the Δ*metE* strain can grow when provided either methionine or B_12_. The amount of B_12_ in a sample is determined by subtracting the growth of the Δ*metE*Δ*metH* control strain from the growth of the Δ*metE* strain after 24 h incubation in the sample and then comparing to a vitamin B_12_ standard curve.

Culture samples and vitamin B_12_ standards were boiled at 100°C for 10 min. Cell debris was removed by centrifugation at 16,000 x*g* for 2 min, and the supernatants were snap frozen in liquid N_2_ and stored at −80°C until analysis. *E. coli* strains were precultivated twice in polystyrene round-bottom tubes in 2 ml M9 medium (Supplementary Data File S2: Media composition) with 0.2% glucose and 1 mg/ml methionine at 37°C and 250 rpm agitation for 24 h. *E. coli* inocula were then collected by centrifugation at 10,000 x*g* for 1 min and washed three times with M9 medium prior to use in the bioassay. Samples and standards were thawed at room temperature and diluted in 2X M9 medium with 0.4% glucose and water to a final volume of 2 ml in polystyrene round-bottom tubes, to which either the Δ*metE* strain or the Δ*metE*Δ*metH* control strain was added to a starting density of 0.01 OD_600_. The bioassay was then incubated at 37°C and agitated at 250 rpm for 24 h, and growth was measured by OD_600_. Sample B_12_ concentration was determined relative to a standard curve.

### Spent medium and cell lysate experiments

Wild-type *C. reinhardtii* was grown for 48 h in continuous light until density reached 1– 4×10^6^ cells/ml. Cells were collected by centrifugation in 500 ml polypropylene bottles for 4 min at 11,900 x*g* and 4°C. The supernatant was passed through a 0.22 µm filter to generate sterile spent medium. Cell pellets were resuspended in fresh medium to the original culture volume and then sonicated using a Fisherbrand Model 505 Sonic Dismembrator (Fisher Scientific, PA, USA) on ice for 45 min with alternating 10 s at 30% intensity and 15 s “off” steps. The supernatant of the resulting cell lysate was then passed through a 0.22 µm filter to generate sterile soluble cell lysate. Spent medium, soluble cell lysate, and parallel live *C. reinhardtii* cultures were then inoculated with *M. japonicum* as described above, and growth was monitored over another 48 h.

To determine the bacterial growth supported by substrates in *C. reinhardtii* cell lysate when these substrates are present at a relevant concentration, wild-type *C. reinhardtii* was grown for 48 h in continuous light, and spent medium and soluble cell lysate were prepared as described above. Then, the NPOC of the two culture fractions was measured, and the cell lysate was diluted to the same NPOC concentration as the spent medium using fresh medium. 3 ml of the diluted cell lysate was inoculated with *M. japonicum* as described above in parallel with spent medium, fresh medium, and fresh medium with 150 µg/ml sucrose in polystyrene round-bottom tubes. Cultures were incubated at 30°C and agitated at 200 rpm, and growth was monitored for roughly 48 h.

## RESULTS

### *M. japonicum* growth is severely limited in coculture with *C. reinhardtii*, especially under diurnal light

Previous work suggested that *C. reinhardtii* supports heterotrophic growth of the rhizobial bacterium *M. japonicum* when cocultured in minimal medium given light and atmospheric CO_2_ [25, 26, 28, 36]. As a first step in characterizing the transfer of fixed carbon in this relationship, we compared the growth of *M. japonicum* when incubated alone or together with wild-type *C. reinhardtii* in minimal medium with either no exogenous carbon source or with 150 µg/ml sucrose (Fig. 1). Coculture with the bacterium did not have a discernable impact on *C. reinhardtii* growth or physiology (Fig. 1B, SI Fig. 2, *p* > 0.05). *M. japonicum* grew considerably with sucrose and only marginally without it, but we found that coculture with *C. reinhardtii* did not significantly increase the maximum bacterial density in either condition (Fig. 1C, *p* > 0.05). Although the maximum density of *M. japonicum* in coculture without sucrose (∼3×10^7^–5×10^8^ CFU/ml) was similar to what has been observed previously [28], we found that the bacterium was able to achieve a comparable density without the alga, in contrast to previous reports.

**Figure 2:**
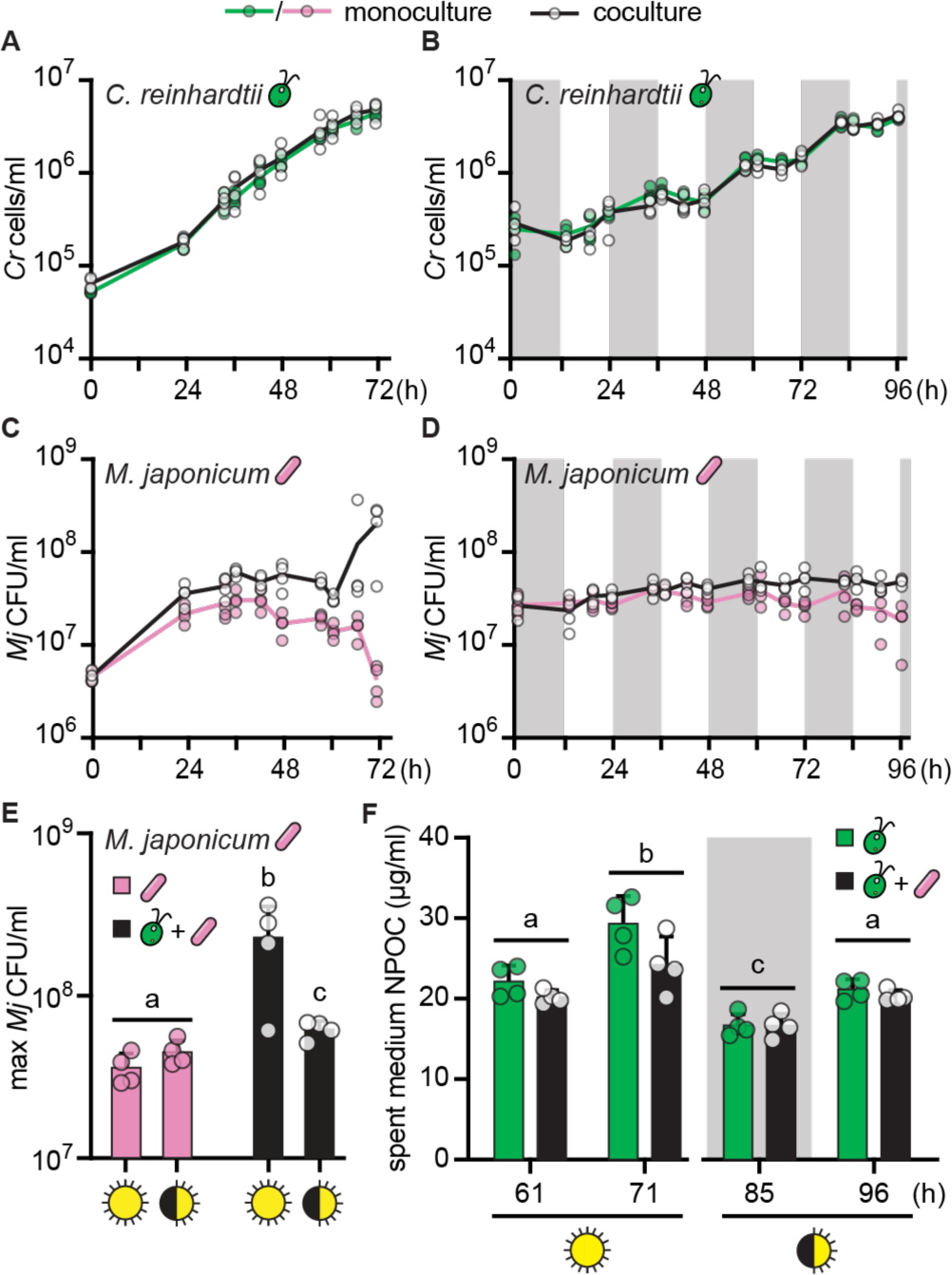
Coculture light regime impacts organic carbon release and *M. japonicum* growth. Quadruplicate (*n* = 4) monocultures (colors) and cocultures (black and white) were bubbled with air and shaken at 125 rpm. Cultures were inoculated to reach the same final algal cell density at the end of the experiment. (A, B) Growth of cell wall reduced strain CC-5390 in continuous light or diurnal light (12-h-light/12-h-dark), respectively, with and without *M. japonicum*. The grey backgrounds indicate sample timepoints that occurred during the dark phase. Under diurnal light, CC-5390 exhibits synchronous growth, doubling roughly once per day upon nightfall. (C, D) *M. japonicum* growth under the continuous and diurnal light regimes, respectively, with and without CC-5390. (E) Maximum bacterial density achieved under the various trophic conditions tested. (F) NPOC in spent medium from CC-5390 and cocultures sampled at the indicated times from the continuous and diurnal light regimes in (A) and (B), respectively. Lower-case letters indicate significantly different groups by paired two-tailed Student’s *t*-test (*p* < 0.05). Error bars represent the standard deviation from the mean.

As the growth of *M. japonicum* was not significantly stimulated by wild-type *C. reinhardtii*, it seemed that the alga did not supply adequate substrates for the bacterium’s heterotrophic metabolism under these conditions. This could be because *C. reinhardtii* released fixed carbon compounds in insufficient quantities, or because *M. japonicum* is unable to metabolize the specific compounds generated by *C. reinhardtii*. We had initially illuminated cocultures with continuous light to maximize algal carbon fixation and growth rate, but we hypothesized that the alga’s nighttime metabolism could stimulate additional growth of *M. japonicum*. During the day, the alga fixes carbon to build biomass and stores photosynthate as starch [10]. After sufficient growth during the light phase, cells initiate DNA synthesis and mitosis in the dark, followed by a G0 phase during which starch reserves are degraded by glycolysis and genes for fermentation are expressed. Fermentation “waste products” released by algae in the night could serve as a readily available source of fixed carbon for cooccurring bacteria.

To test the impact of different light regimes on the interaction, we used the readily synchronized *C. reinhardtii* strain CC-5390. Synchronized populations allow excellent temporal separation of cells in distinct metabolic phases of the cell cycle, and could reveal a diurnal pattern in bacterial growth resulting from the algal partner’s diurnal metabolism. We compared cocultures and monocultures under continuous and diurnal light over several days with high temporal resolution. *C. reinhardtii*’s doubling time is slower under diurnal light (cells only divide once per day), but we designed the experiment to achieve the same final amount of algal biomass under the two light regimes. *C. reinhardtii* strain CC-5390 density reached roughly 4×10^6^ cells/ml under all conditions (Fig. 2A, 2B). This was roughly 2X higher than the density reached by the wild-type strain used in Fig. 1, although the strains appeared to reach stationary phase in the same amount of time under continuous light (72 h). Algal growth was again unaffected by the bacterium. In contrast, we found that *M. japonicum* density increased significantly in the presence of CC-5390 under both light regimes (Fig. 2C–2E). CC-5390 may support more growth of the bacterium than the wild-type strain because it is more prone to cell lysis and releases more nonpurgeable organic carbon (NPOC) into the spent medium (SI Fig. 3). However, we could not discern a clear temporal pattern in bacterial proliferation over the diurnal cycles, and contrary to our hypothesis, we found that continuously illuminated cocultures supported 3-fold more bacterial cells than did diurnally illuminated cocultures (Fig. 2E, *p* = 0.04). Spent medium from continuous light-grown *C. reinhardtii* had significantly more NPOC than spent medium from diurnal light-grown *C. reinhardtii* of the same culture density, suggesting that more dissolved organic carbon was available to the bacterium under continuous light (Fig. 2F, *p* < 0.05). Algal cell lysis was similar under the two light regimes, so differences in algal spent medium NPOC suggest that *C. reinhardtii* exudes more reduced carbon in continuous light than in diurnal light (SI Fig. 4, *p* > 0.05). Finally, the presence of *M. japonicum* did not significantly reduce the concentration of NPOC in the spent medium (Fig. 2F, Student’s *t*-test, *p* > 0.05). This could mean that either *M. japonicum* did not take up significant amounts of organic carbon from the medium or that it could not fully oxidize it to CO_2_, and instead released reduced carbon waste products back into the extracellular milieu.

**Figure 3:**
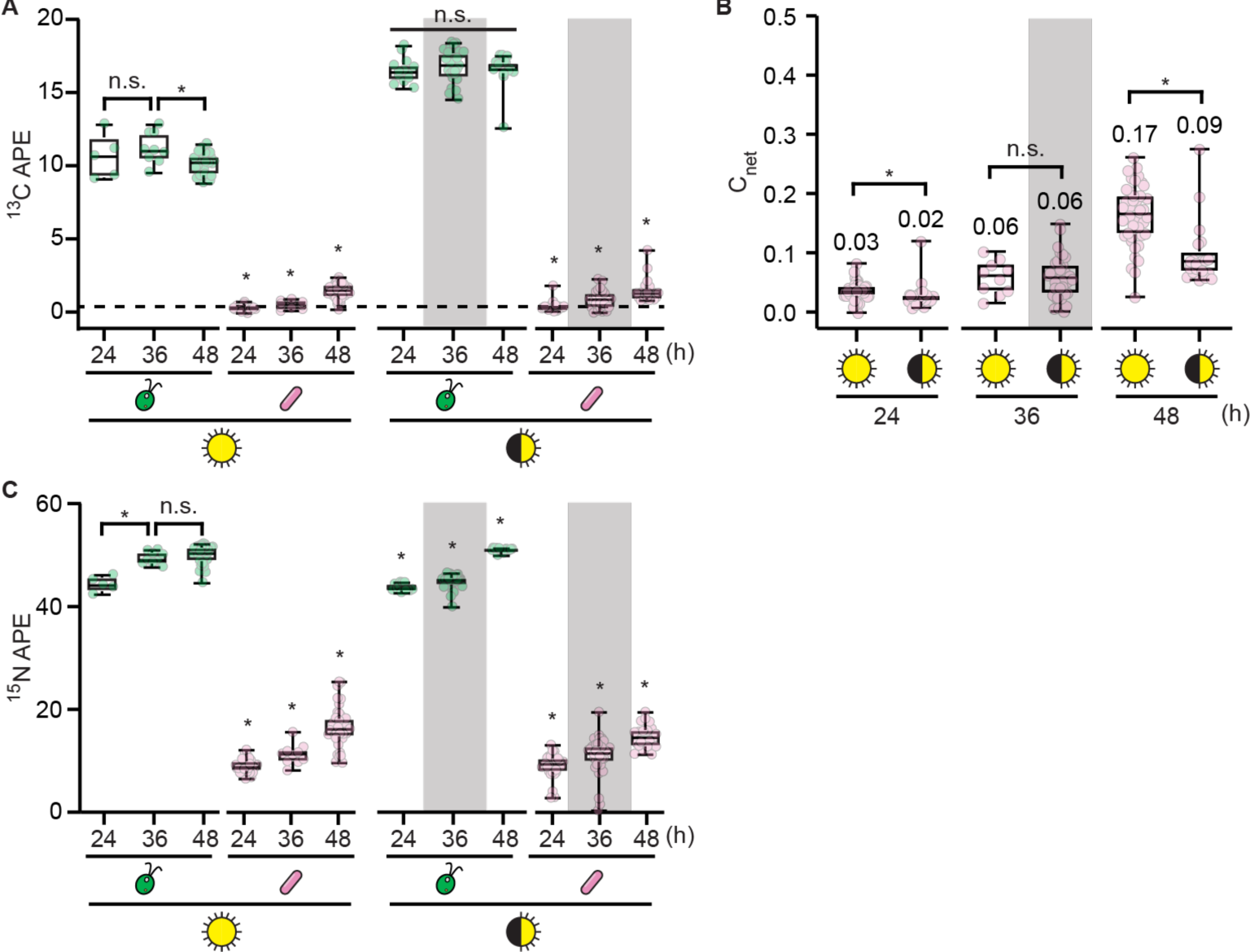
Stable isotope probing demonstrates carbon flux in continuous and diurnal light. Duplicate cocultures (*n* = 2) of CC-5390 and *M. japonicum* were grown as described in Fig. 2 but with 50% of their NH_4_^+^ provided as ^15^N and with ^13^CO_2_ added to the air starting 13 h after inoculation. Isotope enrichment was measured using nanoSIMS at the indicated times after inoculation. The grey backgrounds indicate sample timepoints that occurred during the dark phase. (A) ^13^C atom percent enrichment (APE) data of *n* **≥** 5 individual cells grown under continuous light or diurnal light. The dashed lines represent the maximum ^13^C APE that occurred in *M. japonicum* monoculture controls provided sucrose as a result of nonspecific background or direct CO_2_ incorporation during heterotrophic growth; see SI Fig. 6 for more details. Asterisks indicate significant differences over time in a given culture by two-tailed Student’s *t*-test (*p* < 0.05). (B) C_net_ of *M. japonicum* cells (*n* **≥** 10), calculated relative to the algal biomass as the source. Median values are indicated, and asterisks indicate significant differences by two-tailed Student’s *t*-test (*p* < 0.05). (C) ^15^N APE data of *n* **≥** 5 individual cells grown under continuous light or diurnal light. Asterisks indicate significant differences over time in a given culture by two-tailed Student’s *t*-test (*p* < 0.05).

**Figure 4:**
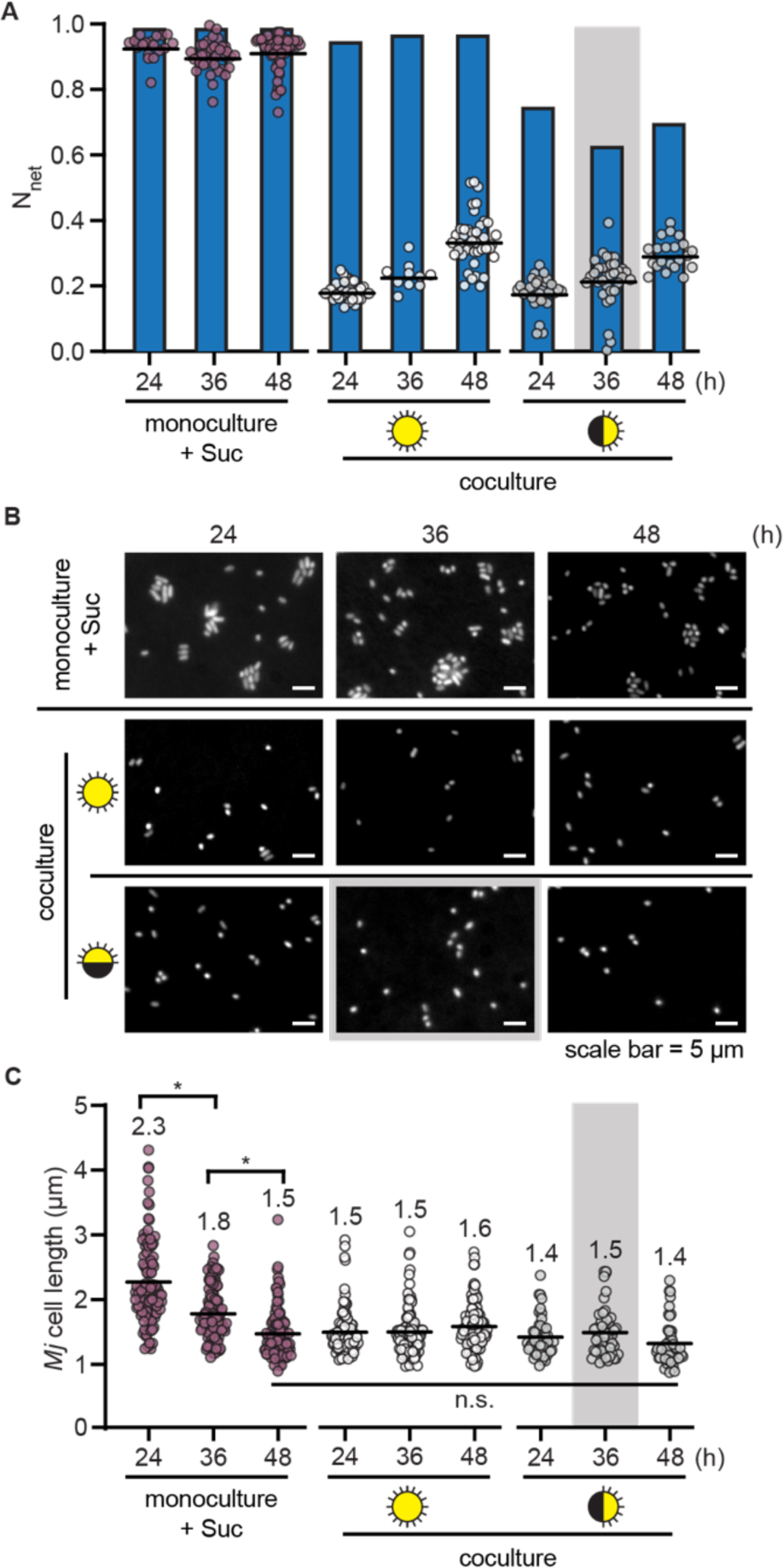
***M. japonicum* undergoes reductive division upon depletion of sucrose and during algal coculture.** Duplicate cultures (*n* = 2) (monoculture with 150 µg/ml sucrose or coculture under continuous light or diurnal light) were grown as described in Fig. 2 but with 50% of their NH_4_^+^ provided as ^15^N and with ^13^CO_2_ added to the air starting 13 h after inoculation. Isotope enrichment was measured using nanoSIMS and cells were imaged by fluorescence microscopy at the indicated times after inoculation. The grey backgrounds indicate sample timepoints that occurred during the dark phase. (A) N_net_ of *M. japonicum* cells (*n* **≥** 10) (circles) relative to the calculated expected N_net_ (blue bars) if each doubling in CFU/ml represented a doubling in biomass (SI Fig. 8). Black lines represent mean values. (B) Representative fluorescence microscopy images of cells stained with DAPI. (C) Cell length of *n* ≥ 50 *M. japonicum* cells per culture type (*n* ≥ 25 cells from each of the duplicate cultures) measured using ImageJ. Mean cell length is indicated, and asterisks indicate significant differences by two-tailed Student’s *t*-test (*p* < 0.05).

### Stable isotope probing reveals mismatch between bacterial biomass synthesis and division in coculture

To determine the amount of carbon fixed by *C. reinhardtii* that is taken up by *M. japonicum*, we conducted stable isotope probing experiments using ^13^CO_2_. Cocultures of CC-5390 and *M. japonicum* grown under continuous and diurnal light as in Fig. 2 were bubbled with air containing ^13^CO_2_. ^15^NH_4_Cl was added to the medium as an independent tracer of biomass synthesis, complementing measurements of cell density. We assayed label uptake with single-cell resolution using nanoSIMS.

We found substantial ^13^C enrichment of the algal cells, which remained at a similar concentration throughout the timepoints examined (Fig. 3A, SI Fig. 5, SI Fig. 6). In addition, we observed ^13^C enrichment of bacterial cells, which increased with time in cocultures under both light regimes (Fig. 3A, SI Fig. 5, SI Fig. 6, *p* < 0.05). Killed controls and monocultures of *M. japonicum* on various substrates demonstrated that this enrichment was not a result of nonspecific background or direct CO_2_ incorporation by the bacterium (i.e. through anapleurotic carboxylation reactions) (SI Fig. 6, SI Fig. 7). Thus, a meaningful amount of carbon derived from algal photoautotrophy was in fact incorporated into *M. japonicum* biomass in the cocultures.

**Figure 5:**
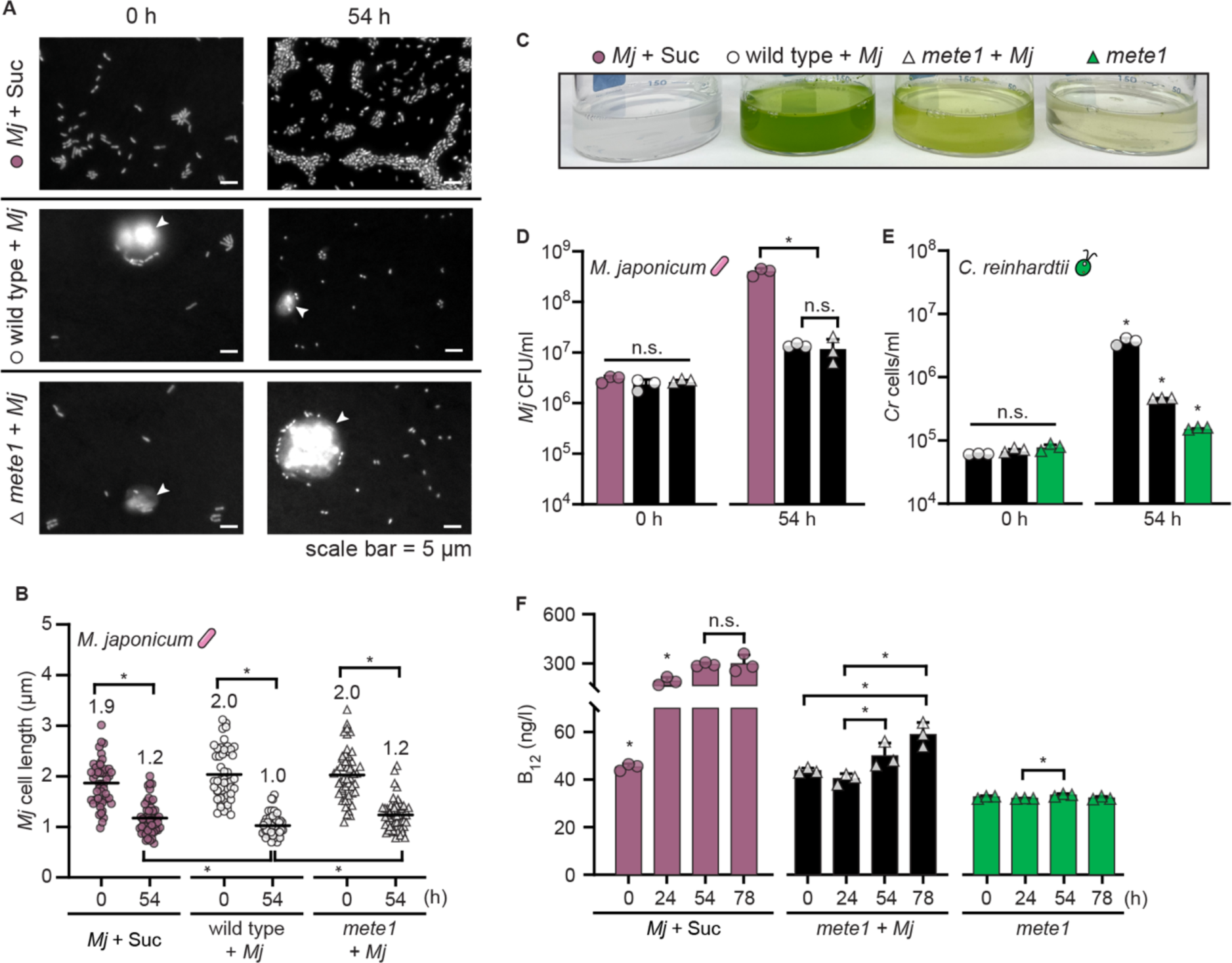
*M. japonicum* delivers vitamin B_12_ to *C. reinhardtii* despite reductive division. Triplicate cultures (*n* = 3) were maintained under continuous light, shaken at 130 rpm, and assayed at the indicated times after inoculation. (A) Representative fluorescence microscopy images of *M. japonicum* with 150 µg/ml sucrose, in coculture with *C. reinhardtii* wild type, and in coculture with *C. reinhardtii mete1*. Cells were stained with DAPI, and *C. reinhardtii* cells are indicated with white arrowheads. (B) Length of *M. japonicum* cells (*n* = 50, 10–20 cells from each of the three triplicate cultures) measured using ImageJ. Mean cell length is indicated. (C) Representative image of cultures 54 h after inoculation. (D) *M. japonicum* density in monoculture with 150 µg/ml sucrose (pink bars) or in coculture (black bars) with wild-type (circles) and *mete1 C. reinhardtii* (triangles), estimated as CFU/ml. (E) Cell density of *C. reinhardtii* wild type (circles) or *mete1* (triangles) in coculture (black bars) and of *mete1* alone without added vitamin B_12_ (green bars). (F) Concentration of vitamin B_12_ in cultures measured using an *E. coli* bioassay. Error bars represent the standard deviation from the mean, and asterisks indicate significant differences by two-tailed Student’s *t*-test (*p* < 0.05).

We compared the extent of carbon transfer under continuous and diurnal light by calculating the fraction of bacterial carbon that was derived from algal carbon, C_net_, using the average ^13^C content of algal biomass over the course of each experiment as a rough measure of the source carbon isotopic composition. We found that significantly more algal ^13^C was transferred to bacterial cells under continuous light over 48 h (0.17 median C_net_ in continuous light, 0.09 median C_net_ in diurnal light) (Fig. 3B, *p* = 6×10^-6^). This was consistent with our observation that spent medium NPOC and *M. japonicum* cell density were greater in cocultures exposed to continuous light than those exposed to diurnal light (Fig. 2, SI Fig. 8A).

Both the ^13^C and ^15^N enrichment in *M. japonicum* cells were lower than we had expected from the increase in bacterial CFUs observed in the experiment (>3-fold increase in CFUs during the first 24 h of coculture under continuous light), suggesting a mismatch between the change in cell density and the bacterial biomass synthesized (Fig. 3C, SI Fig. 8A). We quantified the magnitude of this mismatch using the ^15^N label, for which the source isotopic composition was better constrained, as opposed to that of the ^13^C label which was delivered in the gas phase in an open system. Using the source ^15^N composition and the measured isotopic composition of bacterial cells at each timepoint, we calculated the fraction of newly synthesized biomass generated from the source substrate over the course of the experiment: N_net_ [33]. We compared this to the expected N_net_ that would be observed if each doubling in CFUs in the culture occurred upon a doubling in bacterial biomass (0.99 N_net_ for cells grown on sucrose after 8–13 doublings, 0.63–0.97 N_net_ for cells co-cultivated with *C. reinhardtii* after 1–5 doublings) (SI Fig. 8B). Indeed, the measured N_net_ of *M. japonicum* cells in the cocultures was much lower than the expected values at all measured timepoints (0.18–0.34 N_net_), whereas the measured values matched the expected values during logarithmic-phase growth on sucrose (Fig. 4A). This showed that in coculture with *C. reinhardtii*, *M. japonicum* cells reduced new biomass synthesis by 24 h and yet continued to divide – a hallmark of bacterial starvation often referred to as reductive division [37–39]. As nutrients available for a bacterium’s growth decrease, the cell volume added prior to cell division is reduced, leading to smaller daughter cells [40–43].

If the deviation of the observed N_net_ from the expected N_net_ indeed reflected reductive division, bacterial cells in the coculture would be smaller than cells in the logarithmic phase of growth on sucrose, which are typically 2 µm in length. When we measured the size of the bacterial cells, we found that *M. japonicum* cells were indeed smaller in the cocultures: mean cell length was 1.4–1.5 µm after 24 h in coculture, whereas it was 2.3 µm after 24 h of growth on sucrose (Fig. 4B, 4C). When sucrose was presumably depleted from the medium over the subsequent day in the monoculture, mean cell length decreased to a similar extent, demonstrating that reductive division occurs upon carbon limitation in *M. japonicum*.

Taken together, these data show that photosynthetically fixed carbon was indeed transferred from *C. reinhardtii* to *M. japonicum* under both continuous and diurnal light, and that although this amount was not sufficient for balanced logarithmic-phase growth of the bacterium, it allowed for a small amount of bacterial proliferation.

### Reductively dividing bacteria still deliver sufficient vitamin B_12_ to support growth of the B_12_-requiring *mete1* strain

As reductive division had not previously been reported in the *C. reinhardtii*-*M. japonicum* interaction, we wondered whether the phenomenon also occurs during coculture with the B_12_-requiring *mete1* strain of *C. reinhardtii*, and if so, whether the bacterium still synthesizes adequate B_12_ to support the mutant’s growth. We compared the change in bacterial cell size after growth on 150 µg/ml sucrose to changes during coculture with either wild-type *C. reinhardtii* or *mete1*. In all conditions, we observed a 37–50% reduction in *M. japonicum* cell length over 54 h of cultivation (Fig. 5A, 5B). This suggested that bacterial carbon limitation occurs to a similar extent during coculture with the *mete1* strain as with the wild type.

To test whether reductively dividing *M. japonicum* still delivers vitamin B_12_ to *C. reinhardtii*, we compared the growth of the *C. reinhardtii mete1* strain in coculture and monoculture without exogenous B_12_, and we monitored changes in vitamin B_12_ concentration overtime using a bioassay [34, 35]. *mete1* growth was limited by B_12_ relative to the wild-type *C. reinhardtii*, but the B_12_-requiring strain grew significantly more in the presence of *M. japonicum* despite reductive bacterial division (Fig. 5C, 5E, *p* < 0.05). Furthermore, we found that *M. japonicum* produced significant vitamin B_12_ in coculture, but only after 24 h (Fig. 5F, *p* < 0.05), even though bacterial biomass synthesis and cell size have decreased substantially by that time (Fig. 4). This demonstrates that bacterial B_12_ synthesis and sharing occur despite the carbon limitation experienced during the trophic interaction with the alga.

Although there were roughly 10X more *C. reinhardtii* cells present in the wild-type coculture than the B_12_-limited mutant coculture, there was no difference in *M. japonicum* density between these two conditions (Fig. 5D, *p* >0.05). Thus, greater algal density does not translate into greater *M. japonicum* growth, as noted previously [28]. We observed this phenomenon in cocultures inoculated at several starting relative abundances (SI Fig. 10). This suggests that carbon transfer from the alga to the bacterium could be indirect.

### *C. reinhardtii* lysis can explain observed *M. japonicum* proliferation in coculture

To distinguish the source of the fixed carbon that supports *M. japonicum* proliferation in coculture, we compared *M. japonicum* growth with live *C. reinhardtii* to growth in *C. reinhardtii* spent medium and in the soluble fraction of *C. reinhardtii* cell lysate. Spent medium supported the same number of *M. japonicum* cells as did coculture, demonstrating that the observed increase does not require a direct interaction with the alga (Fig. 6A). Cell lysate from *C. reinhardtii* supported almost an order of magnitude more *M. japonicum* proliferation. Therefore, we hypothesized that the fixed carbon transferred in our cocultures could be sourced from a few lysed *C. reinhardtii* cells rather than from photosynthate exuded by live algal cells.

**Figure 6:**
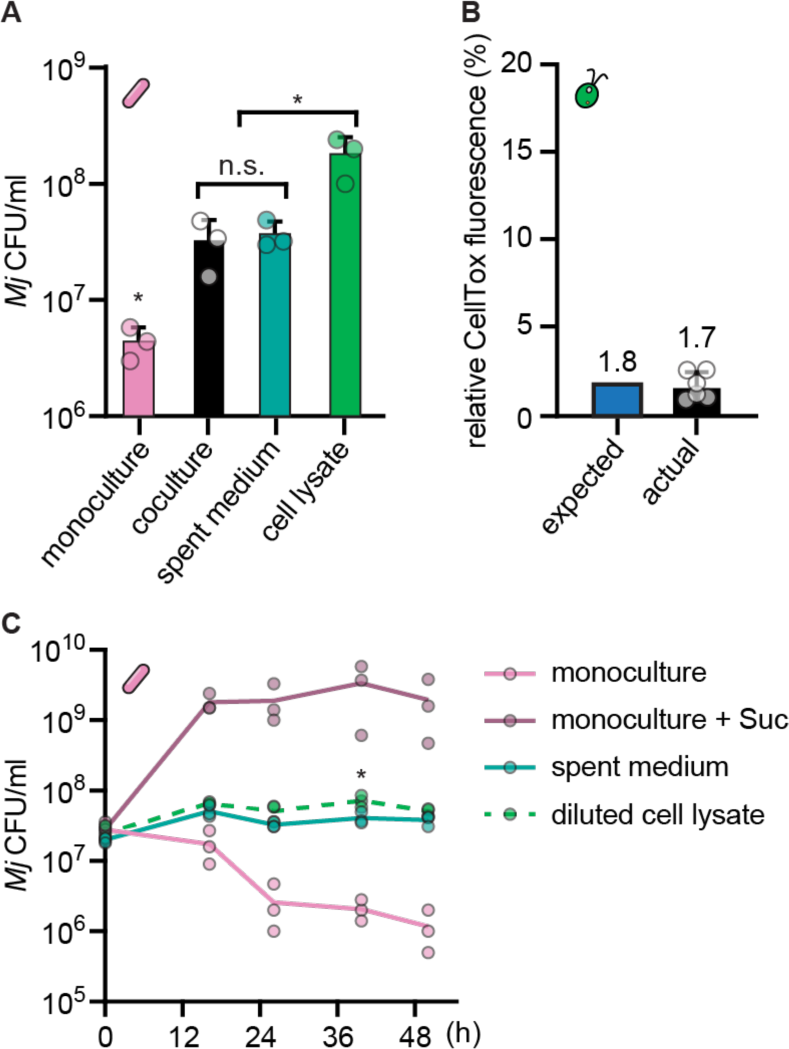
A small degree of algal lysis can explain the *M. japonicum* growth that occurs in coculture. Triplicate cultures (*n* = 3) of wild-type *C. reinhardtii* were grown under continuous light and shaken at 125 rpm for 24 h, reaching 1–4×10^6^ cells/ml. These cultures were then split and used to generate parallel live algal cultures, algal spent media, and algal cell lysate, each then inoculated with *M. japonicum* in triplicate (*n* = 3). (A) *M. japonicum* density 45 h after inoculation in monoculture with no reduced carbon source, coculture with live *C. reinhardtii* cells, *C. reinhardtii* spent medium, or *C. reinhardtii* cell lysate. Asterisks indicate significant differences by two-tailed Student’s *t*-test (*p* < 0.05). (B) The approximate degree of *C. reinhardtii* cell lysis required to explain *M. japonicum* growth in coculture based on *M. japonicum* growth yield in *C. reinhardtii* cell lysate (blue bar), and the degree of *C. reinhardtii* cell lysis measured in the cultures used for the experiment in panel (A) by CellTox Green fluorescence, reported relative to a 100% killed control. The expected value and the mean value are indicated. (C) *M. japonicum* density in triplicate cultures (*n* = 3) over time after inoculation in monoculture with or without 150 µg/ml sucrose, in *C. reinhardtii* spent medium, or in *C. reinhardtii* cell lysate when diluted to the same NPOC concentration as the spent medium. The asterisk indicates a significant difference in density between spent medium and diluted cell lysate 40 h after inoculation by Student’s *t*-test (*p* < 0.05); density was not significantly different between these two conditions at the other timepoints examined (*p* > 0.05). Error bars represent the standard deviation from the mean.

Having determined the increase in viable *M. japonicum* cells supported by lysate from a known density of *C. reinhardtii* cells (1.7×10^8^ CFU/ml on average after 48 h in lysate from 1.6×10^6^ *C. reinhardtii* cells/ml culture), we estimated the theoretical increase in *M. japonicum* CFUs supported by a single lysed *C. reinhardtii* cell to be 100±43 CFU in 48 h. From that, we estimated how many *C. reinhardtii* cells would have to lyse in the coculture to enable the *M. japonicum* proliferation observed: roughly 1.1×10^5^ *C. reinhardtii* cells/ml, which corresponds to 1.8% of the *C. reinhardtii* populations in those cocultures. To test the actual degree of algal lysis in our cultures, we measured algal cellular integrity using the CellTox Green Cytotoxicity Assay. This assay uses a membrane-impermeable DNA dye, resulting in fluorescence from cells whose plasma membrane has been compromised or from extracellular DNA [44]. We compared live culture fluorescence to killed control samples and found that 1–3% of the wild-type *C. reinhardtii* population had compromised cellular integrity (Fig. 6B). Thus, the fraction of dead or dying *C. reinhardtii* cells measured by this assay is comparable to that needed to explain the observed *M. japonicum* proliferation.

Finally, we reasoned that if *M. japonicum* is primarily growing on algal cell lysate, then cell lysate should support the same amount of bacterial proliferation as spent medium when present at the same organic carbon content. Thus, we compared *M. japonicum* growth in spent medium and diluted cell lysate from wild-type *C. reinhardtii* when both were present at roughly 15 µg/ml NPOC. Indeed, algal cell lysate alone supported the same *M. japonicum* growth as spent medium when present at a relevant concentration: bacterial cell density was not significantly different between these two conditions at most timepoints examined (Fig. 6C, *p* > 0.05). The slight increase in *M. japonicum* density in the diluted cell lysate condition could be attributed to increased availability of micronutrients in the fresh medium used to dilute the algal cell lysate. Taken together, these data are consistent with the interpretation that *M. japonicum* growth in coculture is enabled by substrates that are released through *C. reinhardtii* cell lysis rather than by substrates exuded by live *C. reinhardtii* cells.

## DISCUSSION

Most studies of symbioses between *C. reinhardtii* and heterotrophic bacteria to date have focused on impacts on the alga, leaving open the question of how this model phototroph might influence bacterial growth and physiology. Here, we have investigated *C. reinhardtii’s* role as a primary producer in the established coculture system with *M. japonicum*. We have tied carbon cycling in the interaction to alterations in bacterial growth and physiology through quantitative, single-cell analyses. Under continuous light, we found that the alga increased the number of viable bacterial cells, albeit marginally so. We tested whether nighttime metabolism might increase the alga’s propensity for carbon sharing, but we found that diurnal illumination actually decreased the productivity of the partnership. Using nanoSIMS to visualize ^13^C enrichment with high spatial resolution, we quantitatively demonstrated transfer of carbon fixed by the model alga to the bacterium. Roughly 17% of the carbon in *M. japonicum* cells was derived from *C. reinhardtii* over 36 h under continuous light, whereas only ∼9% of the *M. japonicum* carbon came from *C. reinhardtii* over the same amount of time under diurnal light. Bulk measurements showed that algal cultures of similar densities had more organic carbon in their spent medium after continuous illumination than after diurnal illumination. Together, these results suggest that the increased metabolic rate realized under continuous light may be more beneficial for *M. japonicum* (and potentially other bacteria) than fermentative catabolism of starch presumed to occur during the night. It is also possible that although *C. reinhardtii* expresses genes for fermentative metabolism in the night, fermentative metabolites may not actually accumulate to high enough levels in culture unless cells experience anoxia.

With *C. reinhardtii* as the sole source of fixed carbon, *M. japonicum* exhibited signs of starvation, even under continuous light, highlighting the meagerness of the microbial trophic relationship. The quantitative stable isotope probing experiments revealed that during coculture with *C. reinhardtii*, *M. japonicum* synthesized significantly less biomass than expected given the increase in viable bacterial cells. By direct measurement, we showed that *M. japonicum* divided into small daughter cells within 24 h of carbon limitation in coculture, a physiological change often referred to as reductive division. It is well known that as nutrients become limiting, bacterial size at birth and the amount of cell volume each bacterium adds prior to division decrease [40–43]. Studies on freshwater and marine bacteria have documented up to 90% reduction in cell volume during carbon limitation [45–48]. This phenomenon is often considered an adaptive strategy by which diverse bacteria survive starvation [37–39, 49]. By increasing cell number and reducing cell volume, bacteria increase the probability that a member of the population will encounter sufficient nutrients again and increase their surface area-to-volume ratio to improve nutrient uptake. Reductively dividing cells also exhibit changes in stress resistance and macromolecule synthesis. However, we found that despite its apparent starvation, *M. japonicum* provided enough vitamin B_12_ to support a B_12_-dependent *C. reinhardtii* mutant. Vitamin supply enables growth of auxotrophs, influences gene expression and metabolism, and can even give rise to vitamin dependency in algae [25, 50]. Thus, our results suggest that even under austere conditions with meager supply of growth substrates, bacteria can still influence algal physiology and evolution.

We also found that a direct interaction with *C. reinhardtii* was not required to achieve the *M. japonicum* proliferation observed in coculture: growth in algal spent medium yielded the same maximum density of viable *M. japonicum* cells as did coculture. Furthermore, mechanically lysed *C. reinhardtii* cells yielded considerably higher *M. japonicum* densities. This observation led us to hypothesize that perhaps this bacterium does not metabolize the exudates of live *C. reinhardtii* cells at all. Instead, perhaps the small amount of bacterial growth observed in coculture and in spent medium is due to the lysis of a small fraction of algal cells. It has long been appreciated that even during rapid exponential growth, microbial cultures experience a small death rate [51]. Furthermore, cell lysis is an important mechanism of nutrient transfer in natural environments, and microbial necromass is a major component of the soil organic carbon pool [52, 53]. For the experimental conditions examined here, we showed that ∼2% of the *C. reinhardtii* population in our cultures are present as necromass. Based on the *M. japonicum* growth yields achieved in mechanically lysed *C. reinhardtii* cells, we calculated that this is almost exactly the amount of algal necromass required to explain the *M. japonicum* proliferation that occurred in coculture. When present at the same organic carbon concentration, algal cell lysate supported the same amount of *M. japonicum* growth as algal spent medium. Therefore, we propose that lysis is the primary mechanism of trophic transfer in this coculture system, rather than exudation or direct exchange.

Given that very little growth of *M. japonicum* was observed from an interaction with *C. reinhardtii* – even under conditions optimized for algal productivity – our results may suggest that this model alga exhibits a low carrying capacity for heterotrophic bacteria in the environment. Similar bacterial densities have been reported for other *C. reinhardtii* cocultures in minimal media, including those with *Methylobacterium aquaticum* [54, 55]. However, it is possible that other bacteria, such as those that may have co-evolved with *C. reinhardtii* in its native environment, may possess greater capacity to metabolize compounds that *C. reinhardtii* releases during growth. In addition, other bacteria may achieve greater access to growth substrates through chemotaxis, interspecies signaling and direct exchange, or algicidal activity [3, 56–59]. The mode of cell lysis (i.e. mechanical vs. viral) could also greatly impact the algal-derived nutrients released to cooccurring heterotrophs [60]. Nonetheless, stable isotope probing experiments on other algal-bacterial interactions and on seawater microbial communities have reported similar levels of trophic carbon transfer to those we observed here [56, 61, 62]. This suggests that although *C. reinhardtii* and *M. japonicum* may not commonly encounter one another in nature, this relationship may be representative of naturally occurring microbial phototroph-heterotroph interactions. Studying this model system has revealed important aspects of microbial ecology, and our quantitative approach can serve as a roadmap for the future work necessary to understand *C. reinhardtii*’s full potential as a primary producer in the microbial world.

## ACKNOWLEDGMENTS

This work was supported by The Gordon and Betty Moore Foundation Symbiosis in Aquatic Systems Initiative Investigator Award GBML9203 (https://doi.org/10.37807/GBMF9203), and the Lawrence Livermore National Laboratory μBiospheres Scientific Focus Area award SCW1039 from the DOE OBER Genomic Sciences Program. S.D. acknowledges support from NIH T32 Genetic Dissection of Cells and Organisms training grant 1T32GM132022-01. We thank Dr. Alison G. Smith and her group at the University of Cambridge for providing strains and for their generous discussions with us. We thank Dr. Michiko E. Taga at UC Berkeley for her guidance on the work. We thank Dr. Denise Schichnes and the UC Berkeley Biological Imaging Facility for fluorescence microscopy support. We also thank Christina Ramon at Lawrence Livermore National Laboratory and Charles Perrino at UC Berkeley for analytical support.

## DATA AVAILABILITY

All data underlying this article are available in Supplementary Data File S1.

