## Supplementary Information for "Scarcity of fixed carbon transfer in a model microbial phototroph-heterotroph interaction"

**Other supplementary materials for this manuscript include the following:**

Supplementary Data File S1: All data underlying this article (excel file).
Supplementary Data File S2: Media composition (excel file).

### SUPPLEMENTARY METHODS

#### *C. reinhardtii* strain haplotype determination

The haplotype of the wild-type *C. reinhardtii* strain used in this study was determined by allele-specific amplification as previously described [1]. Genomic DNA was extracted using CTAB (hexadecyltrimethylammonium bromide). Cell pellets were resuspended in 0.7 ml 60°C CTAB buffer (2% CTAB, 100 mM Tris-HCl, pH 8, 20 mM EDTA, 1.4 M NaCl, 0.2% beta-mercaptoethanol, 0.1 mg/ml proteinase K) and incubated for 1 h at 60°C. Then, DNA was extracted by addition of 0.7 ml chloroform/isoamylalcohol (24:1), inversion for 2 min, centrifugation at 16,000 x*g* for 10 min at 4°C, and collection of the aqueous phase. DNA was precipitated by overnight incubation in 50% isopropanol at -20°C. DNA was then collected by centrifugation at 16,000 x*g* for 15 min at 4°C, and was then washed with cold 70% ethanol. DNA was air-dried for 15 min at room temperature and resuspended in purified water.

PCR was performed using 300 nM allele-specific primers, 2.5 ng/µl template DNA, and 10 µl iTaq Universal SYBR Green Supermix (Bio-Rad) in 20 µl reactions using a CFX96 Optical Thermocycler (Bio-Rad). Reactions were incubated at 95°C for 30 s followed by 30 cycles of 95°C for 5 s and 63°C for 30 s. Allele-specific primers were the same as previously described [1]. The haplotype of the wild-type *C. reinhardtii* was compared to previously published haplotypes of strains CC-5390 (CC-4351 rescued with the *ARG7* gene), CC-124 (137c *mt–*), and S24- [1].

#### Chlorophyll measurement

Chlorophyll (Chl) content was measured as previously described [2]. 1 ml of culture was collected by centrifugation at 16,000 x*g* for 1 min and the supernatant was discarded. Cell pellets were stored at -80°C until analysis. Pellets were thawed at ambient temperature and then resuspended in 1 ml 80:20 acetone:methanol. Samples were incubated on ice for 5 min, and then starch was collected by centrifugation at 16,000 x*g* for 2 min. Absorbance of the supernatant was measured at 647 nm, 664 nm, and 750 nm using a UV-6300PC Double Beam Spectrophotometer (VWR, PA, USA). As Abs750 was below the detection limit, total Chl was calculated as Chls $a + b$ (µg/ml) = (17.76 x $Abs_{646}$) + (7.34 x $Abs_{663}$) and was then normalized to the *C. reinhardtii* cells in the sample.

#### $F_v/F_m$ measurement

The maximum quantum efficiency of photosystem II ($F_v/F_m$) was measured using a Walz IMAGING-PAM MAXI Chlorophyll Fluorescence System equipped with an IMAG-K7 CCD Camera (Heinz Walz GmbH, Effeltrich, Germany). 300 µl culture was placed in a 96-well plate and dark-adapted for 15 min prior to fluorescence measurement. $F_v/F_m$ was calculated as $(F_m – F_o)/F_m$, where $F_m$ is the maximum fluorescence measured upon a saturating pulse and $F_o$ is the minimal fluorescence of dark-adapted cells in the dark.

### SUPPLEMENTARY FIGURES

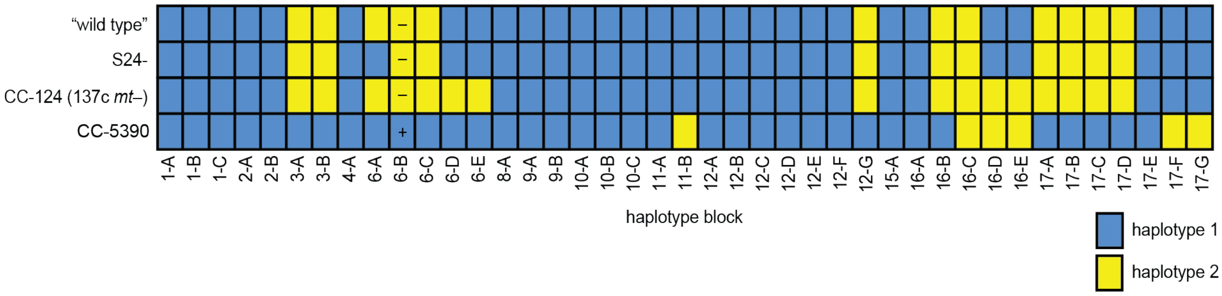


**SI Figure 1: The *C. reinhardtii* "wild type" used in this study is closely related to strain S24-.**

The haplotype of the *C. reinhardtii* "wild type" was determined by allele-specific amplification of genomic DNA and was compared to the reported haplotypes of related strains (S24- and CC-124) and of CC-5390 [1]. Haplotype is indicated at 41 haplotype blocks across the *C. reinhardtii* genome: blue represents haplotype 1, yellow represents haplotype 2, and mating type is indicated as + or – at the mating type locus. Of the *C. reinhardtii* strains for which a haplotype has been reported, the wild-type strain used in this study was most similar to S24-.

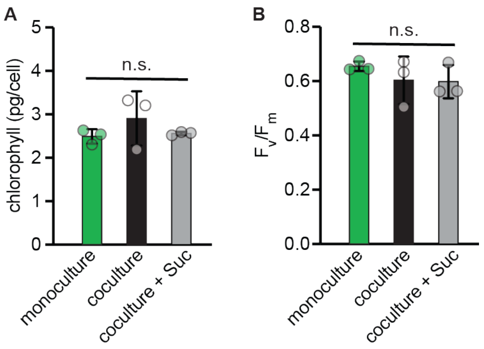


**SI Figure 2: *M. japonicum* does not impact chlorophyll content nor $F_v/F_m$ of wild-type *C. reinhardtii*.**

Triplicate cultures ($n = 3$) were inoculated with or without 150 µg/ml sucrose and maintained under continuous light, bubbled with air, and shaken at 110 rpm. (A) Chlorophyll content normalized to the number of *C. reinhardtii* cells in the cultures 72 h after inoculation. (B) $F_v/F_m$ of cultures 72 h after inoculation. Differences were not significant (n.s.) by two-tailed Student's *t*-test ($p > 0.05$).

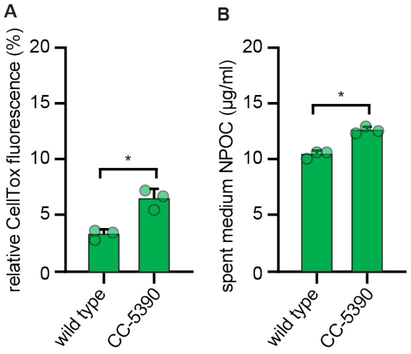


**SI Figure 3: *C. reinhardtii* strain CC-5390 is more prone to cell lysis and release of organic carbon into the medium.**

Triplicate cultures ($n$ = 3) of *C. reinhardtii* wild type and cell wall reduced strain CC-5390 were maintained under continuous light, bubbled with air, and shaken at 125 rpm. (A) The degree of *C.*

*reinhardtii* cell lysis 72 h after inoculation estimated by CellTox Green fluorescence, reported relative to a 100% killed control. (B) NPOC in spent medium sampled 72 h after inoculation. Error bars represent the standard deviation from the mean. Asterisks indicate significant differences by Student's *t*-test ($p < 0.05$).

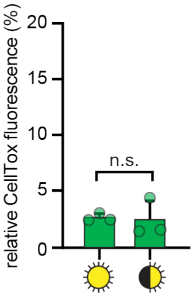


**SI Figure 4: Light regime does not impact *C. reinhardtii* cell lysis.**
Triplicate cultures ($n = 3$) of cell wall reduced strain CC-5390 were maintained in semi-continuous culture in parallel photobioreactors under continuous light or diurnal light (12-h-light/12-h-dark) and bubbled with ambient air. The degree of *C. reinhardtii* cell lysis estimated by CellTox Green fluorescence, reported relative to a 100% killed control, was not significantly different under the two light regimes by two-tailed Student's *t*-tests ($p = 0.86$). Error bars represent the standard deviation from the mean.

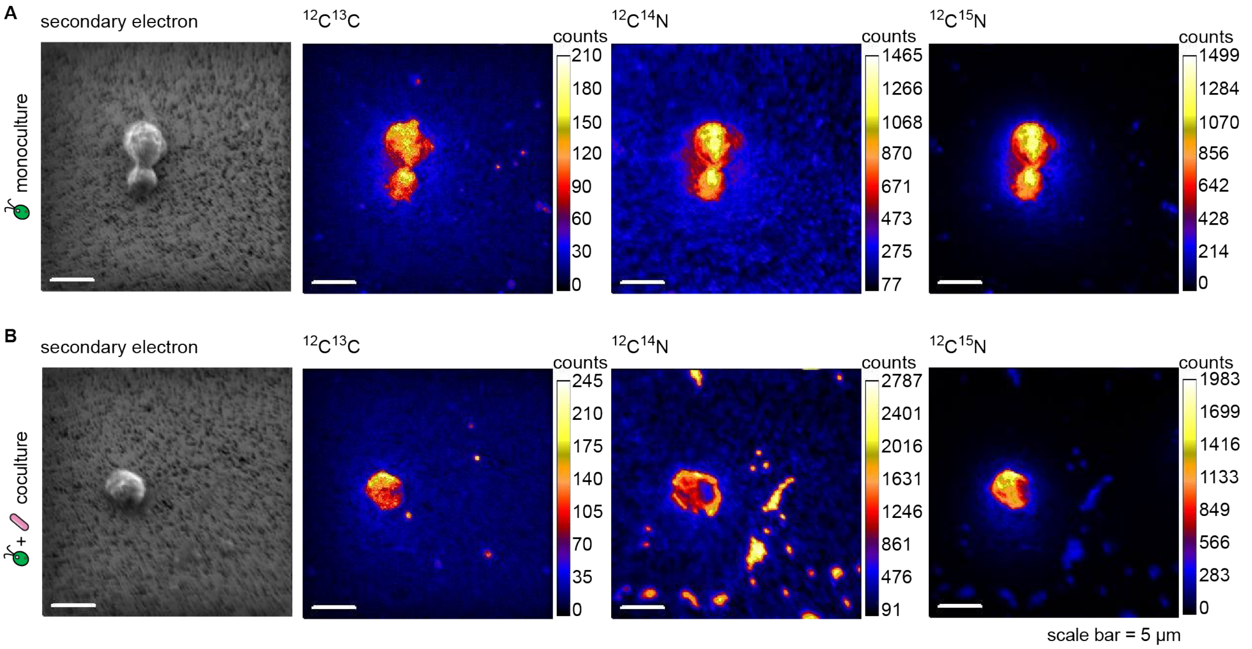


**SI Figure 5: Representative raw nanoSIMS images.**
Representative raw nanoSIMS secondary electron images, $^{12}C^{13}C$, $^{12}C^{14}N$, and $^{12}C^{15}N$ signals from (A) *C. reinhardtii* monoculture and (B) coculture with *M. japonicum*. Algal cells are clearly visible in the scanning electron, $^{12}C^{13}C$, and both CN images. Bacterial cells are most visible in the $^{12}C^{14}N$ image. Micron-scale hotspots in the $^{12}C^{13}C$ image are present in both the monoculture and coculture and do not line up with CN hotspots in the coculture; therefore, they are interpreted as algal debris (e.g., organelles, starch), rather than bacterial cells. Algal ROIs were circled using the

scanning electron and $^{12}C^{13}C$ images and bacterial ROIs were circled using the $^{12}C^{14}N$ image.

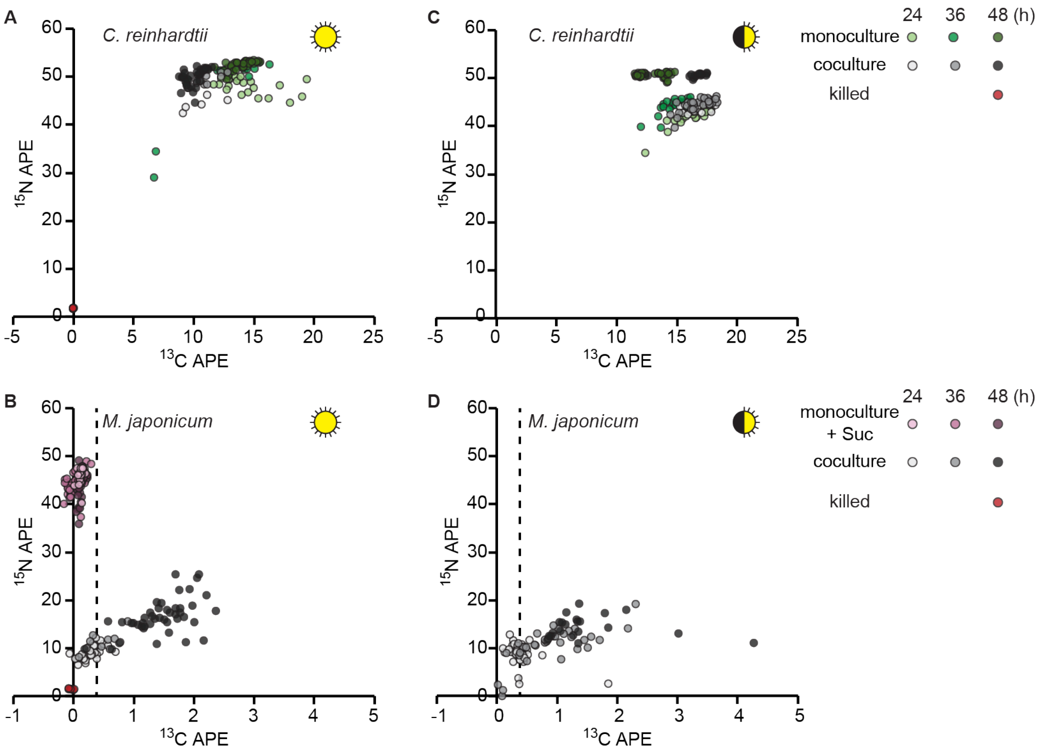


**SI Figure 6: Cross-plots of stable isotope enrichment data.**
Cross-plots of stable isotope enrichment at the indicated times after inoculation in *C. reinhardtii* cells (A, C) and *M. japonicum* cells (B, D), under continuous (A, B) and diurnal (C, D) light, from the coculture experiment described and shown in Fig. 3 plotted alongside monocultures and killed controls. *M. japonicum* grown on sucrose (pink points) is used to constrain possible $^{13}C$ enrichment by direct $^{13}CO_2$ uptake during heterotrophic growth (dashed line), which is well below the maximum $^{13}C$ enrichment observed in cocultures.

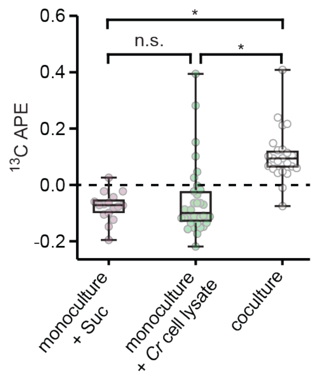


**SI Figure 7: *M. japonicum* $^{13}C$ enrichment is significantly higher during growth with *C. reinhardtii* than during heterotrophic growth on sucrose or unlabeled *C. reinhardtii* cell lysate.**
Duplicate cultures ($n = 2$) of *M. japonicum* were grown with 150 µg/ml sucrose, in *C. reinhardtii* cell lysate, or in coculture with CC-5390 under diurnal light as described in Fig. 3 with 50% of their $NH_4^+$ provided as $^{15}N$ and with $^{13}CO_2$ added to the air used to bubble the cultures starting 13

h after inoculation. Isotope enrichment was measured using nanoSIMS 48 h after inoculation. $^{13}C$ atom percent enrichment (APE) data of $n \geq 16$ individual cells is shown. Asterisks indicate significant differences between the conditions by two-tailed Student's $t$-test ($p < 0.05$).

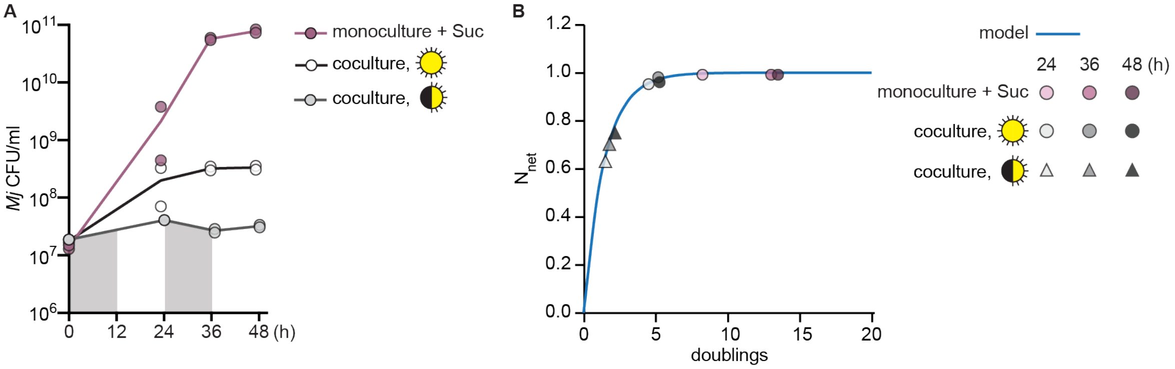


**SI Figure 8: Increase in *M. japonicum* density during stable isotope probing experiments informs $N_{net}$ expected under balanced growth.**

(A) *M. japonicum* cell density during the experiment described in Fig. 3, Fig. 4, SI Fig. 6, and SI Fig. 9 over time after inoculation. The grey background indicates sample timepoints that occurred during the dark phase. (B) Model of $N_{net}$ as biomass doubles (blue) was used to estimate the expected $N_{net}$ of *M. japonicum* cells at each timepoint (circles and triangles) if the number of doublings in CFU/ml observed at that time corresponded to doublings in biomass.

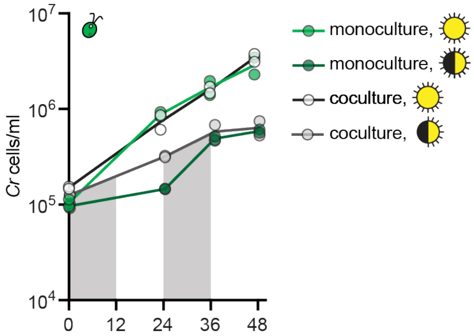


**SI Figure 9: *C. reinhardtii* cell density at the time of nanoSIMS sampling.**

Growth of cell wall reduced strain CC-5390 in continuous light or diurnal light (12-h-light/12-h-dark) with and without *M. japonicum*, during the experiment described in Fig. 3, SI Fig. 6, and SI Fig. 8. The grey backgrounds indicate sample timepoints that occurred during the dark phase.

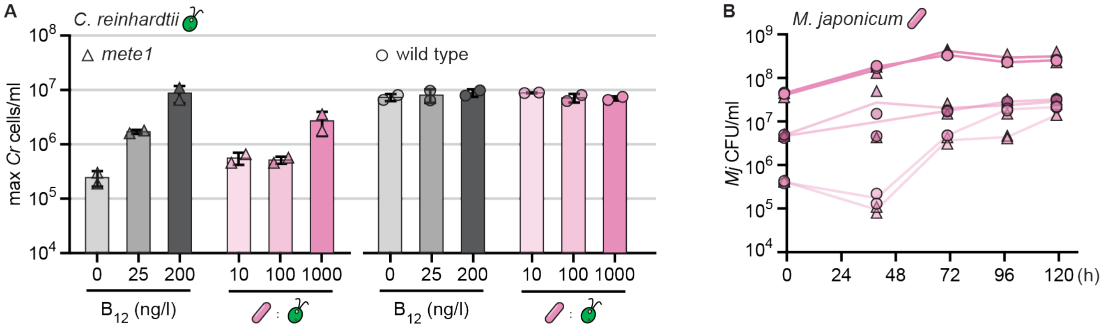


**SI Figure 10: Growth of *C. reinhardtii mete1* strain is supported by vitamin $B_{12}$ and by *M.***

***japonicum*.** Duplicate cultures of *C. reinhardtii* strains with $B_{12}$ or with *M. japonicum* maintained under diurnal light (12-h-light/12-h-dark) and shaking at 130 rpm. (A) Maximum cell density of *C. reinhardtii mete1* mutant (triangles, left) and parental wild-type strain (circles, right) 120 h after inoculation with various amounts of exogenous $B_{12}$ (grey) or with 10, 100, or 1000 bacterial cells per algal cell. Error bars represent the standard deviation from the mean. (B) Growth of *M. japonicum* with *C. reinhardtii mete1* (triangles) or wild type (circles) when inoculated at roughly 10, 100, and 1000 bacterial cells per algal cell (starting density of $5 \times 10^4$ *C. reinhardtii* cells/ml).
